# Evidence for a unique DNA-dependent RNA polymerase in cereal crops

**DOI:** 10.1101/272708

**Authors:** Joshua T. Trujillo, Arun S. Seetharam, Matthew B. Hufford, Mark A. Beilstein, Rebecca A. Mosher

**Author notes:** Corresponding author: Dr. Rebecca Mosher.

## Abstract

Gene duplication is an important driver for the evolution of new genes and protein functions. Duplication of DNA-dependent RNA polymerase (Pol) II subunits within plants led to the emergence of RNA Pol IV and V complexes, each of which possess unique functions necessary for RNA-directed DNA Methylation. Comprehensive identification of Pol V subunit orthologs across the monocot radiation revealed a duplication of the largest two subunits within the grasses (Poaceae), including critical cereal crops. These paralogous Pol subunits display sequence conservation within catalytic domains, but their carboxy terminal domains differ in length and character of the Ago-binding platform, suggesting unique functional interactions. Phylogenetic analysis of the catalytic region indicates positive selection on one paralog following duplication, consistent with retention via neofunctionalization. Positive selection on residue pairs that are predicted to interact between subunits suggests that paralogous subunits have evolved specific assembly partners. Additional Pol subunits as well as Pol-interacting proteins also possess grass-specific paralogs, supporting the hypothesis that a novel Pol complex with distinct function has evolved in the grass family, Poaceae.

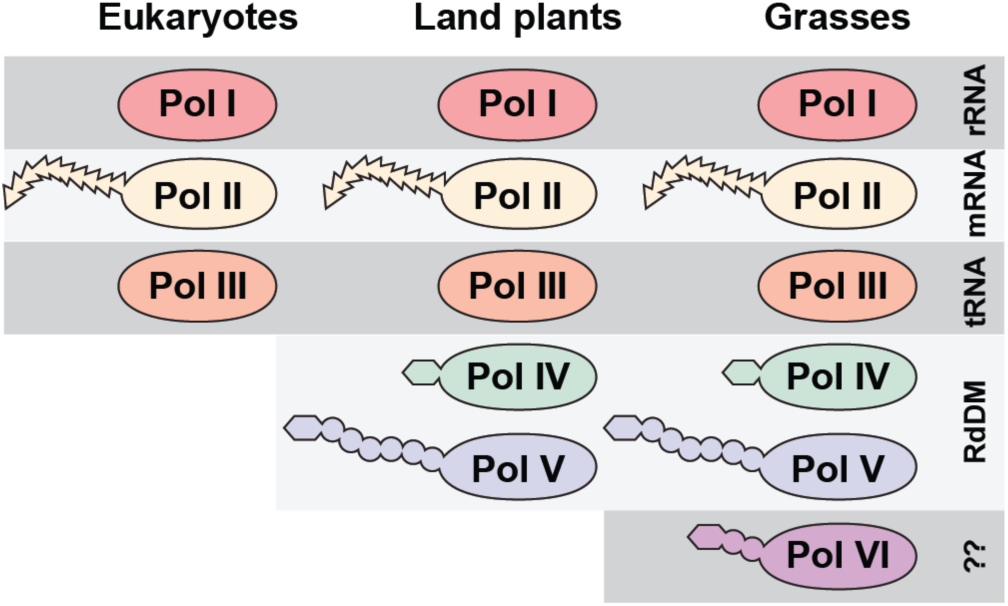
Graphical Abstract.

**Significance statement:** The grass family is critically important for humans, as this group contains cereal grains such as rice, wheat, and corn that form the bulk of the human diet. Here we provide evidence that grasses have evolved a unique polymerase complex of unknown function, suggesting a novel mechanism of gene regulation in the grass lineage. In addition to implications for the biology of grasses, this system offers an opportunity to understand how evolution shapes multi-subunit complexes through duplication of individual components.

## Introduction

Eukaryotic organisms possess three multi-subunit DNA-dependent RNA polymerase complexes (Pol I-III), which are each responsible for transcription of a subset of cellular RNA. Plants encode two additional DNA-dependent RNA polymerases (Pol IV and V), which are specialized for RNA-directed DNA methylation (Haag and Pikaard 2011). RNA Pol IV produces 24-nucleotide small RNAs that are bound by Argonaute4 (AGO4) proteins. These siRNAs then guide AGO4 to sites of Pol V transcription and recruit *de novo* methylation machinery to the locus (Wierzbicki et al. 2009). The carboxy terminal domain (CTD) of Pol V helps to recruit AGO4 through an AGO-binding platform (El-Shami et al. 2007; Lahmy et al. 2016).

DNA-dependent RNA polymerases are composed of multiple subunits, which are named NRPxn, where x=A-E for Pols I-V, respectively, and n=1-12 for the largest to smallest subunit, respectively (Zhou and Law 2015). Pol II, IV, and V share many of their 12 subunits, but also possess unique subunits that confer their distinct functions (Huang et al. 2009; Lahmy et al. 2009; Ream et al. 2009; Haag et al. 2014). The largest subunit of Pol IV (NRPD1) and Pol V (NRPE1) are non-redundant paralogs that together are sister to the largest Pol II subunit (NRPB1) (Luo and Hall 2007). The second, fourth, fifth, and seventh subunits have also duplicated and specialized for Pol IV and V at different times during land plant evolution (Tucker et al. 2010; Huang et al. 2015). In addition, the Argonaute-binding platform in the CTD of NRPE1 is evolving more rapidly than other regions of the protein (Trujillo et al. 2016).

Here, we identify retained duplicates of multiple polymerase subunits and polymerase-associated proteins in Poaceae, the family that contains cereal grasses. Phylogenetic analysis of the two largest subunits indicates positive selection for one paralog following the duplication, consistent with retention via neofunctionalization. Analysis of selection at sites of subunit interaction raises the possibility that evolution favored specific polymerase assemblies. The CTDs of paralogous subunits are also characteristically distinct. Together these results suggest that a sixth distinct RNA polymerase complex exists in this critical plant family.

## Materials and Methods

### Ortholog identification

Published *NRPE1* ortholog sequences (Trujillo et al. 2016) were retrieved from Phytozome versions 11 and 12 (Goodstein et al. 2012). Additional sequences, including homologs of *NRPE1, NRPD1, NRPF1, NRPB2, NRPD/E2, NRPB/D5, NRPE5, NRPB9, AGO4, SPT5/SPT5L*, and *DRD1/CLSY1*-like, were obtained through BLAST or TBLASTX queries against whole genome sequences in Phytozome, CoGE (Lyons and Freeling 2008), and Ensembl Plants (Kersey et al. 2017) using *Oryza sativa* nucleotide sequences. *Streptochaeta angustifolia* and *Zea mays ssp. parviglumis* are available at http://gif-server.biotech.iastate.edu/arnstrm/mhufford/streptochaeta.html and server.biotech.iastate.edu/arnstrm/mhufford/parviglumis.html.

In unannotated genomes, or when gene model predictions were incomplete, coding sequences were predicted using FGENESH+ (Softberry Inc. New York, NY, USA) with *O. sativa* protein sequence as the homology template, followed by manual curation. Orthology was confirmed by reciprocal BLAST searches against the *O. sativa* genome and with phylogenetic analysis. Where species-specific duplications were detected (often due to polyploidization), only one full-length coding sequence was used for downstream analysis. All genes included in this study are listed in Supplemental Table 1.

### Phylogenetic analysis

Nucleotide sequences were aligned by translation using MUSCLE in Geneious version 6.1.8 (Kearse et al. 2012). Where necessary, manual curation was performed to correct alignments. Maximum likelihood phylogenetic trees were inferred with RAxML version 7.2.8 (Stamatakis et al. 2008) using full-length coding sequences for most genes. For the largest subunits (NRPE1 and NRPF1), only the catalytic region from domains B-H were aligned. A General Time Reversible (GTR) model with gamma distributed rate of heterogeneity was implemented, and support values were based on 100 bootstrap replicates.

The branch-sites test for positive selection was performed using PAML version 4.9c codeml (Yang 2007). Branches under positive selection were determined by likelihood ratio test (χ^2^). Parameter space was explored with various starting ω values (0.2, 0.4, 0.6, 0.8, and 1.0) to determine effect on likelihood calculation under M2 (branch-sites) model. Robustness of likelihood values was evaluated by three replications of each analysis under each parameter set.

### Expression analysis

Total nucleic acids were isolated from *O. sativa* inflorescence as previously described (Grover et al. 2017), followed by Turbo DNase-free treatment (Ambion). First-strand cDNA was synthesized using SuperScript III (Invitrogen) with either polyT or random hexamer primers. Ortholog expression was determined through PCR using primers in Supplemental Table 2.

### Structural modeling

Protein homology models of *O. sativa* NRPE1, NRPF1, NRPD/E2, and NRPF2 were generated by Phyre2 intensive modelling (http://www.sbg.bio.ic.ac.uk/∼phyre2) (Kelly et al. 2015) to the *S. pombe* (PDB:3H0G) or *Bos taurus* (PDB:5FLM) Poll II holoenzyme structures for the largest and second largest subunits, respectively. Modelled subunits were then aligned based on interaction of the cognate *S. pombe* subunits and interaction between the subunits was analyzed. Sites under positive selection were visualized in PyMol Molecular Graphics System (Schrödinger, LLC). Residues of *O. sativa* NRPE1 and NRPF1 were compared to homologous residues of NRPB1 in *S. pombe* and NRPE1 in *A. thaliana* with known functional importance. Experimentally derived information regarding subunit interaction regions and specific interacting residues was retrieved through UniProt database (http://www.uniprot.org/) (Bateman et al. 2017).

## Results

### Poaceae members encode paralogous Pol V subunits

Previous studies investigating Pol V evolution revealed the presence of two *NRPE1* paralogs in some monocot genomes (Trujillo et al. 2016). However, this observation was restricted to species within Poaceae as most of the sequenced monocot genomes are members of this agriculturally-important family. To understand the timing of this duplication, we identified putative *NRPE1* homologs across the monocot lineage. Phylogenetic analysis of these sequences reveals a single *NRPE1* ortholog in non-Poaceae monocots and two well-supported clades of *NRPE1*-like sequences within Poaceae (Figure 1A). *Ananus comosus* (pineapple) in Bromeliaceae, a sister lineage to Poaceae in the order Poales, contains a single *NRPE1* ortholog, indicating that duplication of *NRPE1* occurred after the divergence of the two families. Conversely, *Streptochaeta angustifolia*, an early-diverging member of the Poaceae, has two *NRPE1*- like sequences, indicating the duplication occurred in an early ancestor of extant grasses. We have designated the paralog along the longer branch as *NRPF1.*

**Figure 1.**
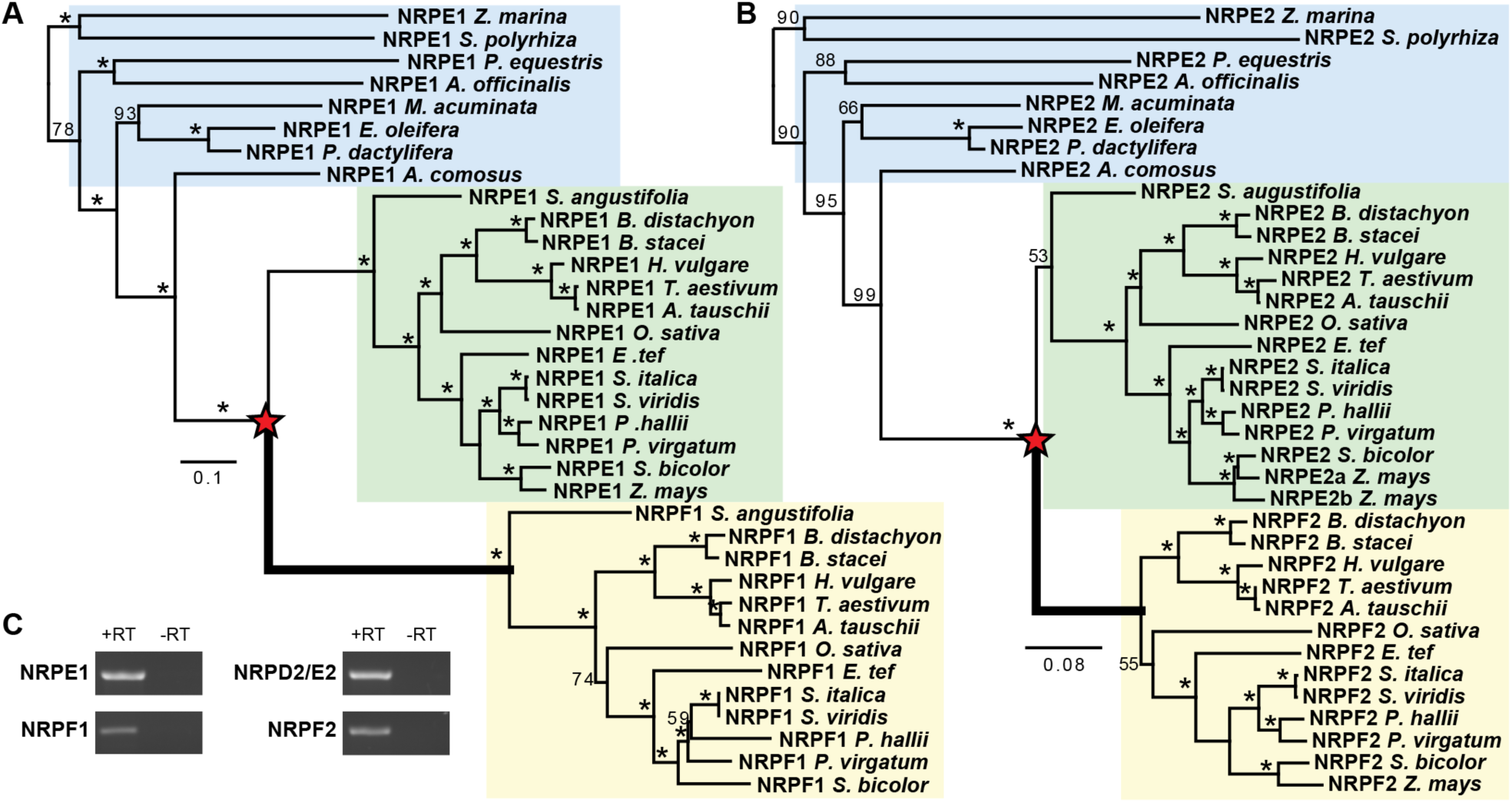
Duplications of Pol V subunits are coincident with the emergence of the grass family Poaceae. The evolutionary relationships of *NRPE1* (A) and *NRPD2/E2* (B) within the monocot lineage demonstrate that monocots outside of the Poaceae family have a single gene copy (blue box), while most members of Poaceae have paralogous genes (green and yellow boxes). Phylogenetic trees were inferred by maximum likelihood analysis of mRNA sequence in the catalytic domain (regions B to H). Bootstrap support values < 100 are shown and red stars mark the inferred duplications. Thick branches indicate positive selection (p < 0.05). Full species names and gene accession numbers are listed in Supplementary Table 1. (C) RT-PCR demonstrates that all *O. sativa* paralogs are expressed.

*NRPE1* and *NRPF1* copies were recovered from every Poaceae genome we assessed, with the exception of *Z. mays*, which lacks a full-length *NRPF1* (Figure 1A). A *NRPF1* homolog is present in the genome, but it appears to be a pseudogene in all *Z. mays* varieties we analyzed (B73, PH207, CML247, and B104), possibly due to insertion of a transposon within the coding sequence (Supplemental Figure 1). Analysis of teosinte (*Zea mays ssp parviglumis*), the wild progenitor of cultivated *Z. mays*, revealed that this pseudogenization occurred prior to domestication. It is not clear why *Z. mays* has lost NRPF1, since retention of this gene in every other grass genome we assessed indicates that these paralogs are not redundant.

In addition to NRPE1, Pol V is distinguished from Pol II by the smaller subunits NRPD2/E2, NRPD4/E4, NRPE5, and NRPD7/E7. We therefore determined if these subunits are also duplicated within Poaceae. We found no evidence of duplication for *NRPD4/E4, NRPE5*, or *NRPD7/E7*, however the second largest subunit, *NRPD2/E2*, which is shared by RNA Pol IV and V, has also duplicated within monocots (Figure 1B).

A single copy of *NRPD2/E2* is present in *A. comosus* but two well-supported clades of Poaceae specific orthologs exist. As with the largest subunit, the clade with the longer branch has been designated as *NRPF2*. It is clear that the *NRPD2/E2* duplication occurred early in Poaceae diversification, however whether this duplication was simultaneous or subsequent to the *NRPE1* duplication is not clear. One *NRPD2/E2* paralog was identified in *S. angustifolia*, and this is sister to all other grass *NRPD2/E2* sequences, suggesting that *NRPF2* might have been lost in *S. angustifolia*. However, it is also possible that *S. angustifolia* diverged from other grasses prior to duplication giving rise to the *NRPD2/E2* and *NRPF2* paralogs since support for the placement of the single *S. angustifolia NRPD2/E2* homolog is weak.

Triplication of *NRPD2/E2* was reported in *Z. mays* (Sidorenko et al. 2009; Stonaker et al. 2009; Haag et al. 2014). With the addition of other Poaceae homologs we show that these three copies arose through a subsequent duplication of *NRPD2/E2* in the maize (or possibly maize + sorghum) lineage, yielding *NRPD2a/E2a, NRPD2b/E2b*, and *NRPF2* (previously called *NRPD2c/E2c*).

We confirmed expression of *NRPE1, NRPF1, NRPD/E2*, and *NRPF2* paralogs in *O. sativa* floral tissue (Figure 1C). Publicly available expression data also support the expression of both paralogs of each subunit (Supplemental Figure 2), confirming that none of the paralogs are pseudogenes. Public expression data also indicate that the paralogs differ in expression level and pattern, supporting our hypothesis that grasses contain novel Pol subunits with non-redundant functions.

**Figure 2.**
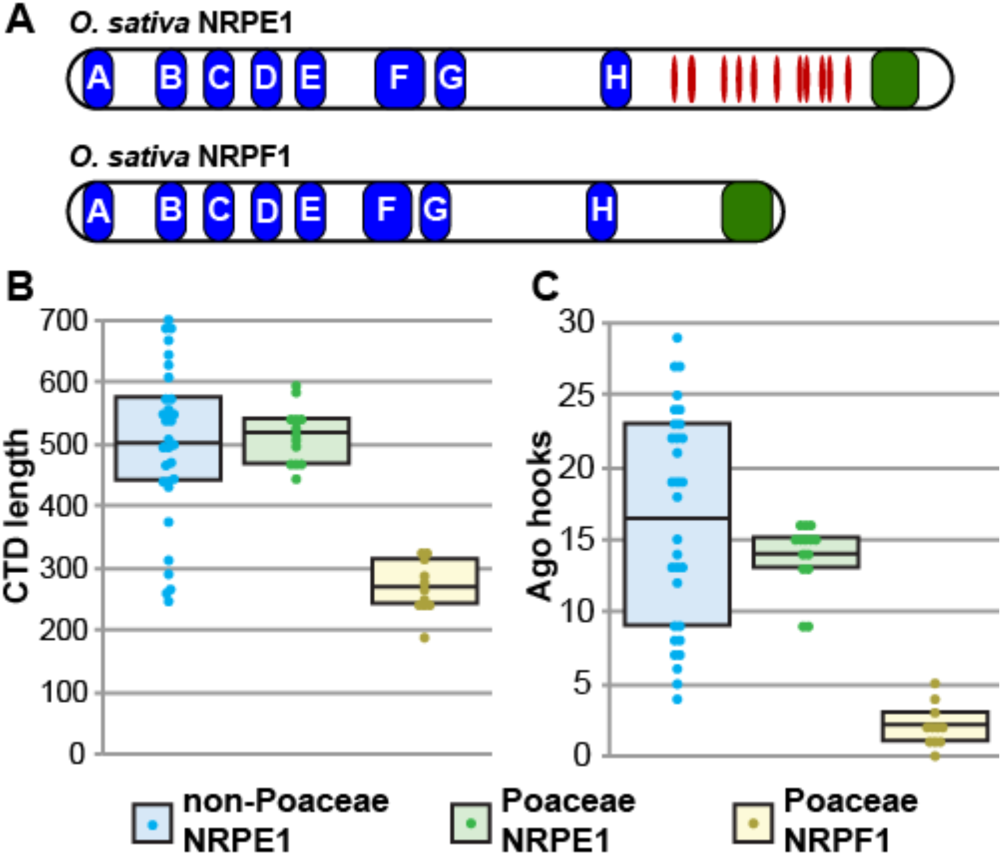
Structural divergence of the CTDs between NRPE1 and NRPF1 paralogs. (A) Diagram of NRPE1 and NRPF1 from *O. sativa*. OsNRPE1 retains a canonical Ago-binding platform between the catalytic A-H domains (blue ovals) and the DeCL domain (green oval). The Ago-binding platform contains many Ago hooks (red ovals). In contrast, OsNRPF1 has a shorter CTD that lacks Ago hook motifs. (B, C) Poaceae NRPE1 and non-Poaceae NRPE1 CTDs are similar in length and number of Ago hooks, while NRPF1 CTDs are shorter and contain fewer Ago hooks. Data points for 31 non-Poaceae, 14 Poaceae NRPE1, and 11 Poaceae NRPF1 are shown as colored circles; boxes represent the interquartile range and the mean is shown by a black bar.

### NRPF1 has distinct CTD characteristics

The largest subunits of Pols II, IV, and V have unique C-terminal domains (CTD). NRPB1 has a large region composed exclusively of heptad repeats; NRPD1 and NRPE1 both contain a domain of unknown function with similarity to Defective Chloroplast and Leaves (DeCL) genes at the extreme C-terminus, but only NRPE1 contains an Ago-binding platform between the catalytic and DeCL domains (Pontier et al. 2005; El-Shami et al. 2007; Huang et al. 2015). The Ago-binding platform is repetitive and disordered, and contains numerous Ago-hook motifs for association with AGO4 (Trujillo et al. 2016). Because the *NRPE1* Ago-binding platform is evolutionarily labile and only closely related sequences can be aligned, CTDs were not included in the phylogenetic analysis that identified *NRPE1* and *NRPF1* clades. However, the same two clades are identified when only characteristics of the CTDs are considered.

NRPE1 CTDs possess an Ago-binding platform similar to that found in NRPE1 from non-Poaceae species, namely a long region with numerous Ago hook motifs. In NRPF1 proteins this region is reduced in length and in number of Ago hook motifs (Figure 2, Supplemental Table 3). However, NRPF1 CTDs are not as short as NRPD1 CTDs, and all but one NRPF1 ortholog contain at least one Ago hook in its CTD, as well as two Ago hook motifs in the catalytic region. This change in CTD character maps to the branch leading to the *NRPF1* clade on the NRPE1/NRPF1 gene tree, suggesting it occurred immediately following duplication. The Ago-binding platform is required for Pol V activity (Wendte et al. 2017), and the change in this domain following duplication further indicates that *NRPF1* is a non-redundant paralog of *NRPE1*.

### Pol VI paralogs experienced positive selection following duplication

Gene duplication allows evolutionary changes that can result in a paralog with a novel function (neofunctionalization) or paralogs that partition the original function (subfunctionalization), hypotheses that can be distinguished based on the pattern of selection following duplication. The branch leading to the *NRPF1* clade is longer than that of the Poaceae *NRPE1* clade, suggesting that *NRPF1* and *NPRE1* experienced different selective pressure following duplication. A longer branch length could result from relaxed selection permitting the accumulation of substitutions; alternatively, positive selection on specific substitutions might have driven the progression towards a novel function. We distinguished between these possibilities with a branch-sites model (Zhang et al. 2005), which indicated that positive selection occurred along the branch leading to the *NRPF1* and *NRPF2* clades while *NRPE1* and *NRPD2/E2* subunits retained purifying selection.

The branch-sites test identified 12 codons with a high likelihood of being under positive selection on the branch leading to *NRPF1* and numerous additional sites when all branches of the clade are evaluated. (Figure 3A, Supplemental Table 4). Similarly, the *NRPF2* branch has 9 sites predicted to be under positive selection immediately following the duplication, and 59 sites across the whole clade (Figure 3B, Supplemental Table 5). Indels also exist between *NRPE1* and *NRPF1* orthologs, suggesting that structural differences were also selected following duplication (Supplemental Figure 3). Only two sites were identified as under positive selection for *NRPE1* and no sites for *NRPD2/E2* (Supplemental Tables 4-5). Positive selection along the *NRPF1* and *NRPF2* branches and predominantly purifying selection for *NRPE1* and *NRPD2/E2* supports the hypothesis of paralog retention due to neofunctionalization, and suggests that in Poaceae, NRPE1 and NRPD2/E2 maintain the ancestral functions, while NRPF1 and NRPF2 may have evolved novel functions.

**Figure 3.**
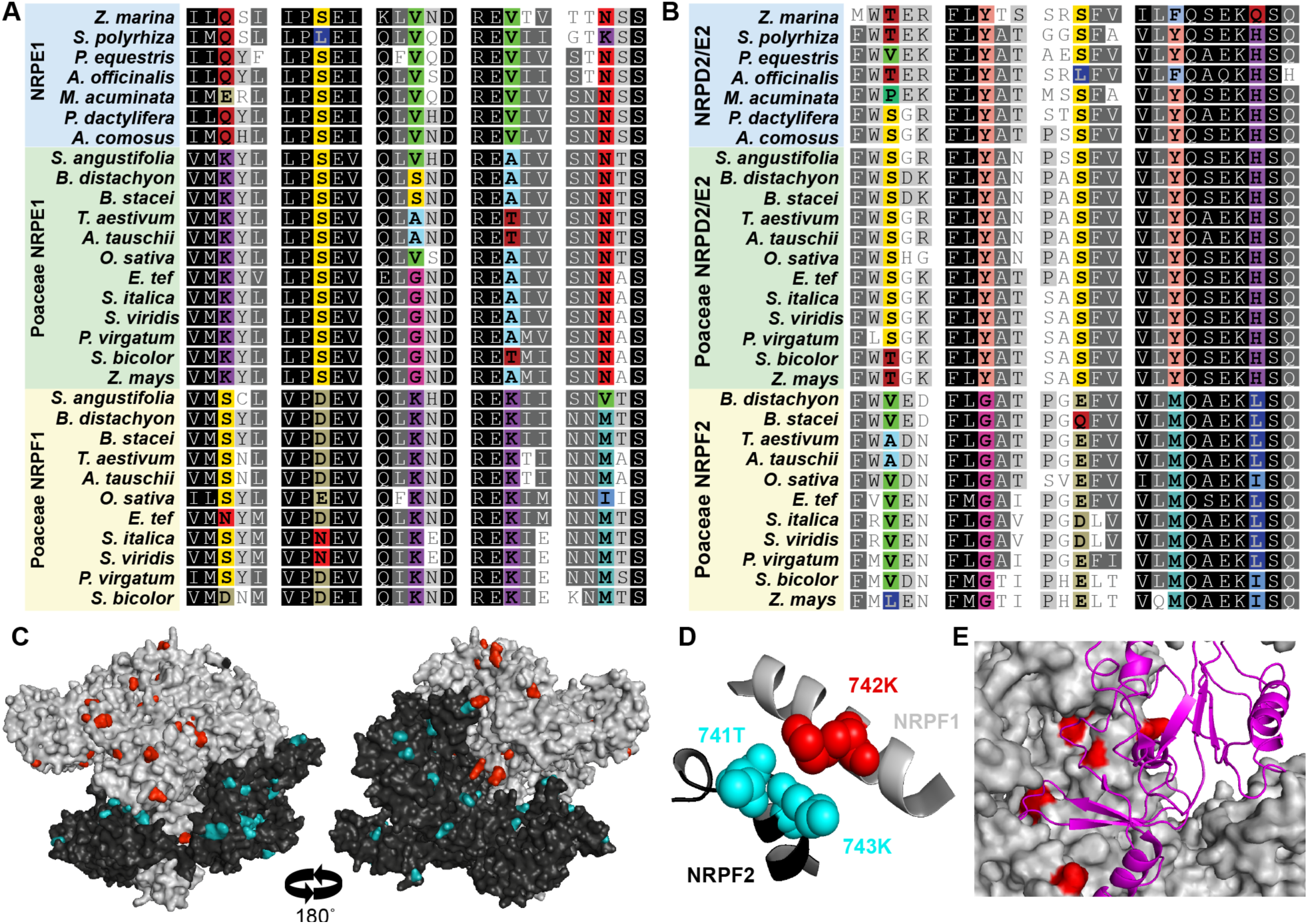
Sites under positive selection cluster on the surface of Pol VI subunits. Alignment of monocot largest (A) and second largest (B) subunits illustrates residues that are under positive selection following gene duplication (colored) (A). Remaining residues are colored based on sequence conservation. (C) Residues under positive selection (colored) are found on the surface of homology-based structures of NRPF1 (gray) and NRPF2 (black). (D) Residues under positive selection also occur at the interface between subunits, as demonstrated by 742K in NRPF1, which is directly opposite 741T and 743K in NRPF2. (E) Several residues under positive selection are found where the largest subunit (gray) interacts with the fifth subunit (magenta ribbon).

### Sites under positive selection are exposed and predict additional subunit duplications

We mapped the predicted sites under positive selection on a homology-based model of *O. sativa* paralogs to evaluate the structural significance of particular substitutions (Figure 3C). NRPE1 and NRPF1 were aligned and modelled to the largest subunit of *S. pombe* RNA pol II holoenzyme (PDB:3H0G chain A) with 100% confidence and sequence identity of 23%. The second largest subunit paralogs were analyzed in the same manner with *O. sativa* NRPD2/E2 and NRPF2 mapping with 100% confidence and 36-37% sequence identity to a bovine RNA Pol II structure (PDB:5FLM chain B).

Most sites predicted to be under positive selection were located on the surface of subunits or at interaction faces with other polymerase subunits, suggesting that the substitutions do not impact the overall structure of the subunits, but might impact assembly of the holoenzyme (Supplemental Tables 4-5). In one case, residues under selection in NRPF1 and NRPF2 are predicted to interact, suggesting that they might be compensatory, and selection may have acted to restrict the number of possible holoenzyme assemblies (Figure 3D). Specific assembly of Pol subunits would indicate that not only are the paralogous subunits functionally non-redundant, they assemble into a unique polymerase complex, a Pol VI.

Based on the position of selected sites on the structure model, we hypothesized additional duplications of smaller subunits. Six selected sites are in the region that interacts with the fifth subunit (Figure 3E) and four sites near the interaction region for the ninth subunit (Supplemental Tables 4-5). Although there is no evidence for duplication of the Pol V-specific *NRPE5*, we identified multiple copies of *NRPB5/D5* in Poaceae (Supplemental Figure 4). There is also evidence for a duplication of *NRPB9/D9/E9* within the monocots (Supplemental Figure 5), although this duplication is difficult to resolve given the limited sequence information in this small subunit. Duplication of *NRPB5/D5* suggests that a Poaceae-specific Pol VI might have assembled in a modular fashion, using paralogous modules/subunits from both Pol IV and Pol V.

**Figure 4.**
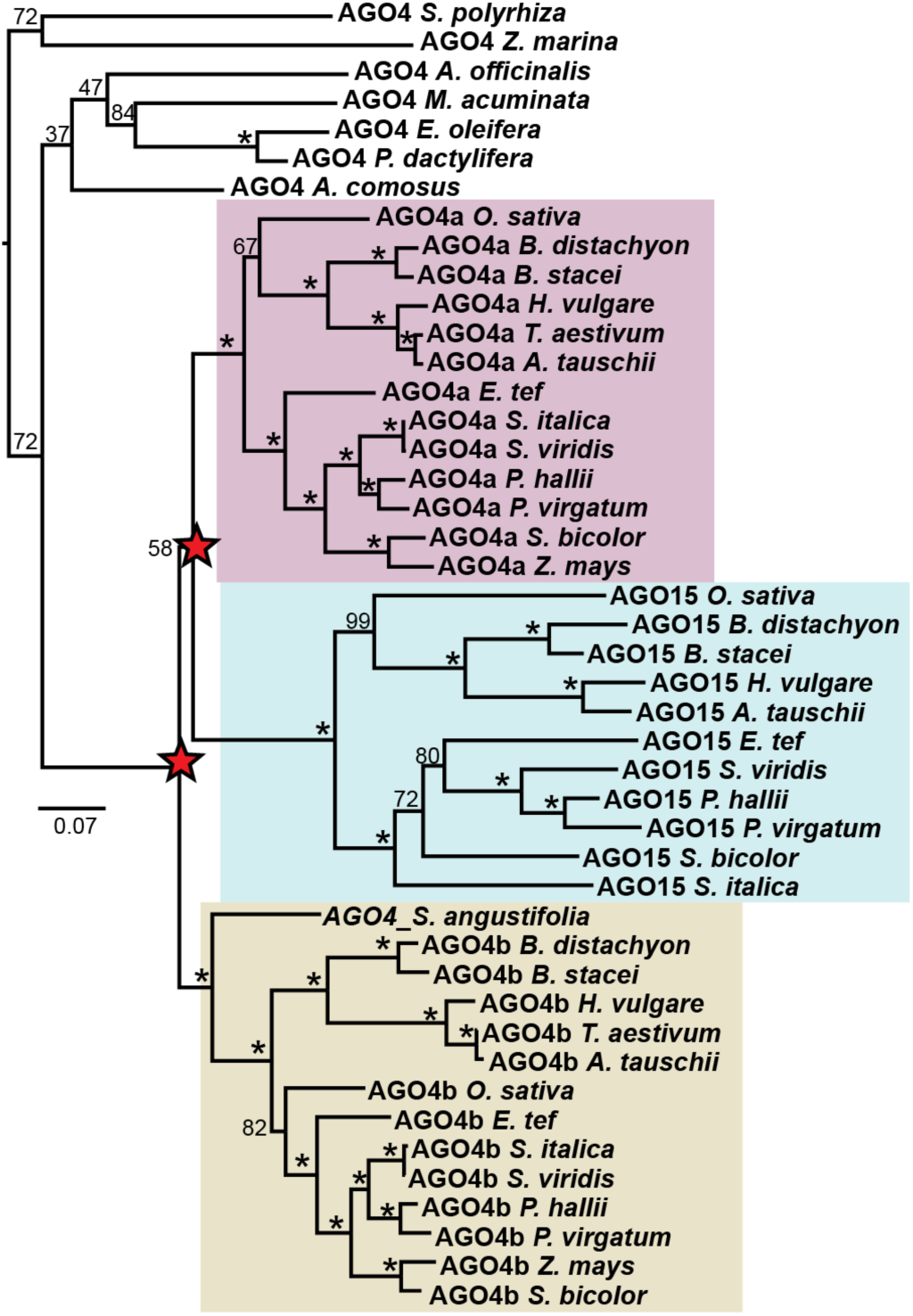
Nested duplications of AGO4 locus result in three paralogs in most grasses. Comparison of *AGO4*-orthologous sequences in monocots demonstrates that grasses contain multiple *AGO4* paralogs and that these duplications were coincident with the emergence of the Poaceae family. Maximum likelihood phylogenetic tree of *AGO4* related nucleotide sequences in monocots. Support values for branches with <100 bootstrap support are marked. Red stars mark the inferred duplication events.

**Figure 5.**
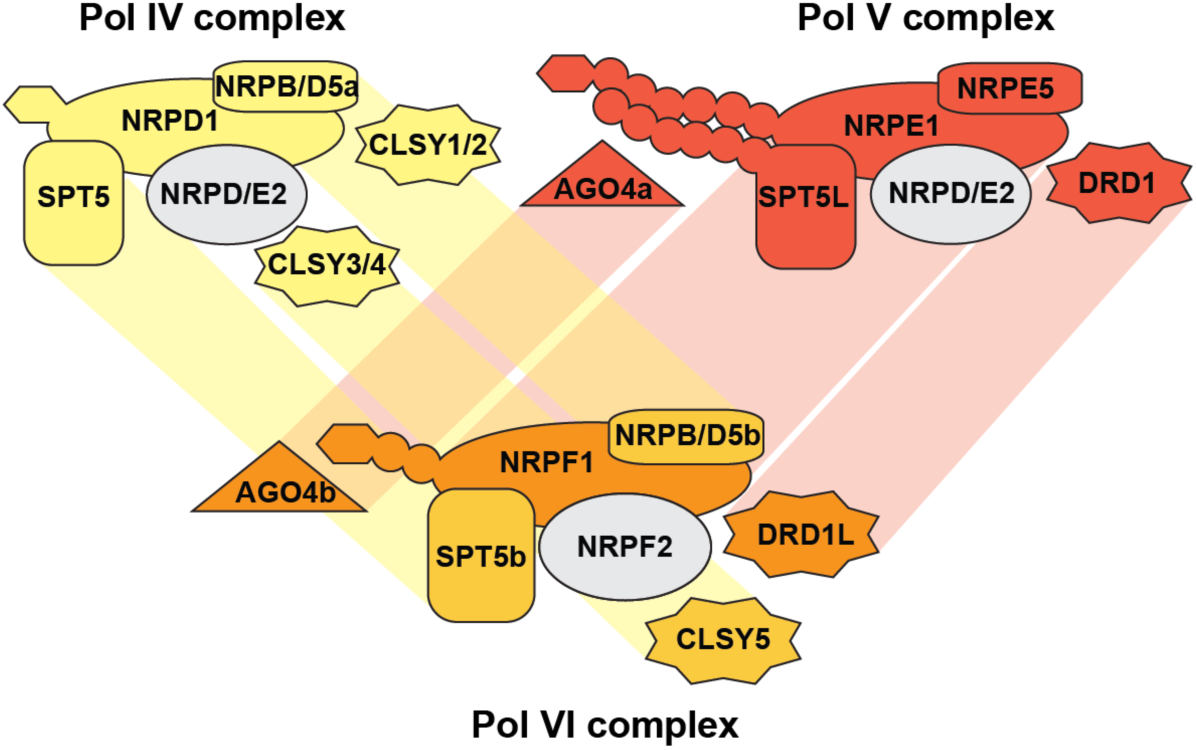
Proposed Pol assembly in grasses. We propose that NRPF1 and NRPF2 subunits assemble into Pol VI with a paralog of NRPB/D5. The Pol VI complex might function with specific paralogs of SPT5, DRD1, CLSY3/4, and AGO4, highlighting the use of paralogous proteins from both Pol V and Pol IV complexes. Paralogous subunits are connected by shaded boxes.

### Pol V-associated proteins are also duplicated in Poaceae

Canonical RdDM involves RNA Pol V interacting with numerous other proteins to accomplish DNA methylation. If Pol VI has a novel function that is diverged from Pol V, we might expect duplication and neofunctionalization of Pol V interacting proteins. We therefore investigated the evolution of known Pol V-interacting proteins, including the small RNA-binding protein Argonaute 4 (AGO4) (El-Shami et al. 2007), the transcriptional elongation factor SPT5-like (Huang et al. 2009), and the SWI/SNF-related helicase DRD1 (Law et al. 2010; Zhong et al. 2012).

AGO4 associates with the NRPE1 CTD, enabling base-pairing of AGO4-bound small RNAs with Pol V transcripts (Wierzbicki et al. 2009), or with the Pol V transcription bubble (Lahmy et al. 2016). In *A. thaliana, AGO4* is a part of a group of Argonautes including the deeply-conserved *AGO6*, and Brassicaceae-specific *AGO8* and *AGO9* (Zhang et al. 2015; Rodríguez-Leal et al. 2016). In addition to an *AGO6* group, we detected three well-supported clades of AGO4 orthologs within grasses arising from nested duplications at the base of Poaceae (Figure 4). *AGO4a* and *AGO15* are sister groups that share high sequence similarity and are located within a few kilobases of one another, suggesting that a tandem duplication of one paralog occurred following whole genome duplication. Most predicted *AGO15* sequences consist of partial, fragmented coding sequences, and there is no evidence for *AGO15* protein accumulation in rice (Wu et al. 2010), suggesting that *OsAGO15* might be an expressed pseudogene. Public expression data indicate that rice and maize AGO4 orthologs are broadly expressed (Supplemental Figure 6), where they bind to different groups of small RNAs (Wu et al. 2010). Genetic data in maize also indicates that despite the fact that *ZmAGO4a* (*ZmAGO119*) and *ZmAGO4b* (*ZmAGO104*) have broad and overlapping expression patterns, these paralogs are not redundant (Singh et al. 2011).

SPT5L, a duplicate of the Pol II transcription elongation factor SPT5, interacts with Pol V and contains its own Ago-binding platform in its carboxy terminus (Bies-Etheve et al. 2009; Lahmy et al. 2016). Although SPT5L is the paralog that interacts with Pol V, we do not detect a duplication of this gene. Rather, *SPT5*, which interacts with Pol II and Pol IV, has undergone a duplication at the base of Poaceae (Supplemental Figure 7). This observation is additional evidence that only specific components of Pol V were duplicated in grasses, and further supports the hypothesis that a sixth polymerase complex formed through duplication of both Pol IV and Pol V machinery.

RNA Pol IV and V transcription is assisted through interaction with helicase proteins of the DRD1-like family (Kanno et al. 2004; Smith et al. 2007; Law et al. 2011). We discovered Poaceae-specific duplications within the *DRD1* and *CLSY3/4* clades, giving rise to paralogs we have named *DRD1-like* and *CLSY5*, respectively (Supplemental Figure 8). We discovered *DRD1L* and *CLSY5* copies only in Poaceae species, though *DRD1* and CLSY trees suggest the duplication predates the evolution of the grasses. We take the current placement as a preliminary assessment in need of more data from additional species to more fully resolve these gene tree topologies. Whether one or both helicases are required for transcription by a grass-specific Pol VI remains to be determined.

DRD1 interacts with RDM1 and DMS3 to form the DDR complex, which is required for RdDM (Law et al. 2010). We identified only a single copy of *RDM1* and *DMS3* in Poaceae, further demonstrating that many components of Pol V machinery remain in single copy, while specific components of Pol VI and Pol V duplicated in grasses. The duplication of Pol IV and Pol V interacting protein further supports our hypothesis that grasses contain a distinct sixth polymerase with unique activity.

## Discussion

Our evolutionary analysis of DNA-dependent RNA polymerases within the monocot lineage of land plants identified duplications of multiple subunits and polymerase-associated proteins. These duplications are coincident with the *rho* whole genome duplication at the base of grasses (McKain et al. 2016). Most genes return to single copy following whole genome duplication, therefore retention of duplicated genes is evidence for the formation of non-redundant protein function (Hahn 2009). Verified expression of *NRPE1, NRPF1, NRPD2/E2*, and *NRPF2* (Figure 1C); unique CTD sequences for *NRPE1* and *NRPF1* (Figure 2); and phylogenetic evidence of positive selection on *NRPF1* and *NRPF2* (Figure 3AB) indicate that these paralogous subunits are not merely redundant, but rather are likely to encode unique functions.

Many of the sites with evidence for positive selection are at interfaces where Pol subunits interact (Figure 3DE), suggesting that in addition to selection for unique activity, there might have been selection for faithful assembly of subunits into unique complexes (Beilstein et al. 2015). This idea is supported by biochemical evidence from *Z. mays*, in which NRPD/E2 and NRPF2 display differential association with NRPD1 (Haag et al. 2014). However, pseudogenization of *NRPF1* in *Z. mays* makes this species a poor representative for other grasses and further validation of Pol subunit associations is required in a different grass species. The signature of selection at predicted interacting sites, as well as the coordinated duplication of multiple subunits, leads us to hypothesize that not only do *NRPF1* and *NRPF2* encode novel functions, but that they assemble into a unique polymerase complex, Pol VI.

We identified duplications of many Pol V subunits and interacting proteins, but we also found duplications of proteins that are not specific to Pol V. For example, we detected duplication of *NRPB/D5*, but not *NRPE5* (Figure S3). Similarly, the Pol V-specific transcription elongation factor *SPT5L* is not duplicated, but the paralogous *SPT5*, which interacts with Pol II and Pol IV, occurs in multiple copies (Figure S6). Duplication of Pol II/IV proteins suggest that Pol VI formed through neofunctionalization of both Pol IV and Pol V subunits (Figure 5). Pol IV and Pol V both generate non-coding transcripts from otherwise silent DNA, but they differ in their speed, accuracy, and processivity (Wierzbicki et al. 2008; Zhai, Bischof, et al. 2015; Marasco et al. 2017). Conservation of key enzymatic residues, including the metal binding sites, indicates that Pol VI is capable of transcription, although such activity and how it differs from Pol IV and Pol V remain to be studied.

One key difference between Pol IV and Pol V is association with AGO4. The NRPE1 CTD contains numerous Ago-hooks for interaction with AGO4. Likewise, it’s binding partner SPT5L also contains numerous Ago hooks and associates with AGO4 (Bies-Etheve et al. 2009; Trujillo et al. 2016). In contrast, neither NRPD1 nor SPT5 contain Ago-hook motifs for AGO4 interaction. NRPF1 orthologs possess only a few Ago hooks, suggesting that Pol VI might interact with AGO proteins, but in a manner distinct from the Pol V-AGO4 interaction. Duplication of *SPT5* rather than *SPT5L* also hints that numerous Ago hooks are not necessary for Pol VI function.

The biological role of Pol VI remains to be determined, but its similarity to Pol IV and Pol V along with the presence of Ago hooks suggests a role in small RNA biology while expression data indicate that Pol VI might accumulate during reproductive development (Supplemental Figure 2). Pol VI might be required for the biosynthesis or function of a number of novel small RNA classes, including highly expressed endosperm-specific siRNA loci in rice (Rodrigues et al. 2013), 24-nt phased siRNAs required for microspore development (Zhai, Zhang, et al. 2015; Fei et al. 2016), or rice “long” miRNAs (Wu et al. 2010). The potential role of Pol VI in reproductive development could make it an important target of agricultural and biotechnology manipulation.

Grasses are one of the most successful radiations of land plants, covering vast areas of natural habitat and agricultural land and forming the bulk of the human diet. We are acutely dependent on grasses, both for food and environmental stability, and it is therefore critical to understand the unique gene regulatory mechanisms of this family. Our discovery of a novel sixth polymerase in Poaceae uncovers a previously unknown aspect of grasses and offers an opportunity to learn more about this important plant family.

## Acknowledgements

The authors thank Dr. May Khanna and Dr. Samantha Perez-Miller for assistance with visualization of homology models. We also thank Dr. Elizabeth Kellogg for support in the assembly of *S. angustifolia* and *Z. mays ssp parviglumis* genomes and Dr. Lynn Clark for providing *S. angustifolia* tissue. We gratefully acknowledge support from the National Science Foundation (IOS-1546825 to R.A.M. and M.A.B) and the National Institutes of Health (T32-GM008659 to J.T.T).

